# The effects of drought and nutrient addition on soil organisms vary across taxonomic groups, but are constant across seasons

**DOI:** 10.1101/348359

**Authors:** Julia Siebert, Marie Sünnemann, Harald Auge, Sigrid Berger, Simone Cesarz, Marcel Ciobanu, Nathaly R. Guerrero-Ramírez, Nico Eisenhauer

## Abstract

Anthropogenic global change alters the activity and functional composition of soil communities that are responsible for crucial ecosystem functions and services. Two of the most pervasive global change drivers are drought and nutrient enrichment. However, the responses of soil organisms to interacting global change drivers remain widely unknown. We tested the interactive effects of extreme drought and fertilization on soil biota ranging from microbes to invertebrates across seasons. We expected drought to reduce the activity of soil organisms and fertilization to induce positive bottom-up effects *via* increased plant productivity. Furthermore, we hypothesized fertilization to reinforce drought effects through enhanced plant growth, resulting in even dryer soil conditions. Our results revealed that drought had detrimental effects on soil invertebrate feeding activity and simplified nematode community structure, whereas soil microbial activity and biomass were unaffected. Microbial biomass increased in response to fertilization, whereas invertebrate feeding activity substantially declined. Notably, these effects were consistent across seasons. The dissimilar responses suggest that soil biota differ vastly in their vulnerability to global change drivers. As decomposition and nutrient cycling are driven by the interdependent concurrence of microbial and faunal activity, this may imply far-reaching consequences for crucial ecosystem processes in a changing world.

## Introduction

Anthropogenic global environmental change affects ecosystem properties worldwide and threatens important ecosystem functions^1,2^. Climate change is predicted to alter precipitation regimes towards more frequent and severe drought events in the future^3^. Simultaneously, human activities, such as fossil fuel combustion and fertilization, are causing an acceleration of the turnover rates of the nitrogen cycle and will double nitrogen deposition in the future^4,5^. The same is true for phosphorous inputs, which also increased at a global scale^6^. Thus, multiple global change drivers are occurring side by side, and their effects are not necessarily additive or antagonistic. Our knowledge on their interactive effects, however, is still highly limited^7,8^. This is particularly true for the responses of soil organisms, which mediate crucial ecosystem functions and services, such as nutrient cycling and decomposition^9,10^. Their significant role is not adequately reflected in the body of global change literature yet. Therefore, a more comprehensive understanding of above- and belowground dynamics is key to predict the responses of terrestrial ecosystems in a changing world^7^.

Many soil organisms are dependent on a water-saturated atmosphere or on water films on soil aggregates^11-14^. Altered precipitation patterns will result in drought periods, which are likely to have substantial effects on their abundances and community structure, thus affecting important soil organism-mediated ecosystem processes. Previous studies reported detrimental effects of drought on soil microbial respiration and biomass as well as a reduction of the diversity of microbial communities^15^. Furthermore, drought was shown to cause a decline in soil microarthropod abundances^16^. In contrast, drought seems to have only marginal effects on nematode community composition^17^. Yet, a reduction of soil moisture content can induce community shifts *via* lower trophic levels, often favouring fungal-feeding nematodes over bacteria-feeders, as fungi perform relatively better under dry conditions^17,18^.

Nutrient enrichment is another key factor that affects the soil community by altering the physical and chemical properties of the soil, e.g., by influencing pH-value, soil porosity, and organic fractions^19-21^. Nitrogen addition has been identified to decrease soil microbial respiration and biomass, often leading to shifts in the soil microbial community composition under the use of mineral fertilizer (NPK)^22-24^. On the other hand, fertilization treatments were shown to increase soil microbial catabolic and functional diversity^25,26^. Furthermore, nitrogen addition alters the nematode community structure towards bacterivores, thus promoting the bacterial-dominated decomposition pathway^27^, and was shown to simplify communities^17^. At the same time, nitrogen enrichment is one of the major drivers determining aboveground primary production^28^. Nitrogen and phosphorous addition are known to increase total aboveground biomass and consequently the quantity and quality of plant litter input to the soil^26,29^. This enhances resource availability *via* bottom-up effects and can therefore increase soil microarthropod abundances^30^. Concurrently, the fertilization-induced increase in aboveground biomass may cause higher transpiration rates, which are likely to reinforce drought effects on soil organisms^31^.

To investigate the interactive effects of extreme drought events and fertilization (NPK), we established a field experiment at the UFZ Experimental Research Station (Bad Lauchstädt, Germany), which combines the treatments of two globally distributed networks - the Drought-Network and the Nutrient Network^32^. Here, we tested the responses of soil microorganisms, nematodes, and soil mesofauna to the interactive effects of extreme drought and fertilization (NPK) across all seasons. Based on prior research, we hypothesized that (1) drought will reduce the activity of soil organisms, whereas (2) fertilization will increase their activity, owing to enhanced plant litter input that subsequently increases resource availability for soil organisms. Furthermore, we predicted that (3) the interactive effects of drought and fertilization will result in detrimental conditions for soil organisms as the negative effects of drought were expected to be further enhanced by increased plant growth under fertilization, resulting in reduced soil water availability for soil organisms.

## Methods

### i. Research site

The study site is located at the Experimental Research Station of the Helmholtz Centre for Environmental Research (UFZ), which is situated in Bad Lauchstädt, Germany. The field site is located in the central German dry area with a mean annual precipitation of 487 mm and an average annual daily temperature of 8.9°C (Meteorological data of Bad Lauchstädt, Helmholtz Centre for Environmental Research GmbH - UFZ, Department of Soil System Science, 1896-2017). The area represents an anthropogenic grassland, which is maintained by moderate mowing (twice a year since 2012). It is a successional plant community dominated by *Vulpia myuros* (L.) C. C. Gmel., *Picris hieracoides* (L.) and *Taraxacum officinale* (F. H. Wigg.) with *Apera spica-venti* (L.) P. Beauv. and *Cirsium arvense* (L.) Scop. being very common. The soil is classified as a haplic chernozem, developed upon carbonatic loess substrates, distinguished by a composition of 70% silt and 20% clay^33^.

### ii. Weather conditions

Weather conditions within the two-year sampling period of this study were in line with the long-term average despite some exceptions: precipitation patterns deviated from the long-term average in 2016 with a dry May (21.2 mm compared to 62.3 mm of the long-term record from 2005-2015) and a wet June (80.2 mm compared to 41.2 mm of the long-term record from 2005-2015). September tended to be dryer than usual in both years (19.5 mm in 2016 and 22.1 mm in 2017 compared to 51.8 mm of the long-term record from 2005-2015).

### iii. Experimental design and treatments

The experimental site was established in March 2015. The experimental design consists of five blocks with five plots each. The plots have a size of 2 × 2 m and are arranged at a distance of 3 m from each other (Fig. S1). The experiment includes two treatments with two levels each (first applied in March 2016): drought (control/drought) and fertilization (no NPK/NPK addition), as well as their interaction (drought x fertilization). Notably, this experiment crosses treatments of two globally distributed experimental networks: the full NPK fertilization treatment of the Nutrient Network^32^ and the drought treatment of the Drought-Network (http://www.drought-net.colostate.edu/).

In order to simulate drought, a rainfall manipulation system was established^34^ using corrugated acrylic strips. The roofs have a size of 3 × 3 m and reduce precipitation by 55% throughout the year, simulating a severe long-term reduction in precipitation. Roofs were built with a slope of 20° to ensure water runoff and account for the expected snow load in the region. Exclusion of potential artefacts was realized by equal roof constructions using inverted acrylic strips conceived to let rainfall pass^35^ (Fig. S2). To control for possible infrastructure effects of the roof constructions itself, a fifth plot was added to each block without any roof construction (ambient plots), thus receiving ambient precipitation (not crossed with the fertilization treatment and thus not part of this study, see Fig. S1). To validate the drought treatment, soil water content was quantified on all plots in every sampling campaign. All three precipitation levels differed significantly in their soil water content (Tukey’s HSD test, p < 0.05): as intended, the lowest soil water content was found for the drought treatment (−19.4% compared to the ambient plots). Also the infrastructure control plots (with concave roof constructions) differed significantly from the ambient plots (without roof construction), indicating that there were effects of the roof construction itself (−13.4%). Furthermore, soil water content varied significantly between seasons (Table S1; Fig. S3).

The fertilization treatment was realized by annual addition of a mixture of nitrogen (N), phosphorus (P) and potassium (K) (i.e. NPK fertilization; applied at 10 g m^−2^ y^−1^ by elemental mass) before each growing season. In addition, the micronutrient mix “Micromax Premium” (Everris) was applied in the first treatment year^32^.

### iv. Soil sampling

The first soil sampling took place in March 2016. Sampling campaigns were repeated every three months to cover every season (spring, summer, fall, winter) from March 2016 to December 2017 (i.e., eight samplings across two years). Samples were taken on all plots with roof construction (drought and control) with a steel core sampler (1 cm in diameter; 15 cm deep). Seven subsamples per plot were homogenized, sieved at 2 mm, and stored at 4°C. Soil samples were used to determine soil water content and microbial respiration. In addition, nematodes were extracted from the soil samples in spring and summer of 2017.

### v. The Bait Lamina Test

Feeding activity of soil invertebrates was surveyed using the bait lamina test (Terra Protecta GmbH, Berlin, Germany), which presents a commonly used rapid ecosystem function assessment method^36^. The test uses rigid PVC sticks (1 mm × 6 mm × 120 mm) with 16 holes of 1.5 mm diameter in 5 mm distance. Original sticks were filled with a bait substrate consisting of 70% cellulose powder, 27% wheat bran, and 3% activated carbon, which was prepared according to the recommendations of Terra Protecta. The bait substrate is primarily consumed by mites, collembolans, nematodes, enchytraeids, millipedes, and earthworms, whereas microbial activity plays a minor role in bait loss^37-40^. The sticks were inserted vertically into the soil with the topmost hole just below the ground surface. To avoid damaging the sticks, a steel knife was used to prepare the ground prior to insertion. Five sticks were used per plot to account for spatial heterogeneity^41^. For each sampling campaign, the bait lamina sticks were removed from the soil after three weeks of exposure and evaluated directly in the field. Bait consumption was recorded as empty (1), partly empty (0.5), or filled (0). Thus, soil invertebrate feeding activity could range from 0 to 16 (maximum feeding activity). Mean bait consumption per plot was calculated prior to statistical analyses.

### vi. Microbial biomass and activity

An O2-microcompensation system was used to measure the respiratory response of soil microorganisms^42^. First, basal respiration was determined as a measure of soil microbial activity (μl; O_2_ h^−1^ g^−1^ soil dry weight). Second, the maximal respiratory response after the addition of glucose (4 mg g^−1^ dry weight soil, solved in 1.5 ml distilled water) allowed us to determine microbial biomass μg Cmic g^−1^ soil dry weight)^43^.

### vii. Nematode analysis

Nematode extraction was conducted with a modified Baermann method^44^. Approximately 25 g of soil per plot were transferred to plastic vessels with a milk filter and a fine gaze (200 μm) at the bottom and placed in water-filled funnels. More water was added to saturate the soil samples and to ensure a connected water column throughout the sample and the funnel. Hence, nematodes migrated from the soil through the milk filter and the gaze into the water column and gravitationally-settled at the bottom of a closed tube connected to the funnel. After 72 h at 20°C, the nematodes were transferred to a 4% formaldehyde solution. Nematodes were counted at 100x magnification using a Leica DMI 4000B light microscope. Identification was conducted at 400x magnification. For identification, sediment material from the bottom of each sample vial was extracted with a 2 ml plastic pipette and examined in temporary mounted microscope slides. At least 100 well-preserved specimens (if available in the sample) were randomly selected and identified to genus (adults and most of the juveniles) or family level (juveniles), following Bongers (1988)^45^. Nematode taxa were then arranged into trophic groups (bacteria-, fungal- and plant-feeders, omnivores and predators)^46,47^. Due to low densities, omnivorous and predatory nematodes were grouped into a combined feeding type for most analyses. Nematodes were also ordered according to the colonization-persistence gradient (c-p values)^48,49^. The colonizer-persistence scale classifies nematode taxa based on their life history strategy (i.e. r or K strategists). Cp-1 taxa are distinguished by their short generation cycles and high fecundity. They mainly feed on bacteria. Cp-2 taxa have longer generation times, lower fecundity and consist of bacterivores and fungivores^50^. Both are categorized as r-strategists. Cp-3 to cp-5 are classified as K-strategist nematodes with longer generation times, higher trophic feeding levels and increasing sensitivity against disturbances^50^. The c-p-values can be used to calculate the Maturity Index (MI) as weighted means of nematode families assigned to c-p-values. It is used to describe soil health and as an indicator of overall food web complexity^48,49^.

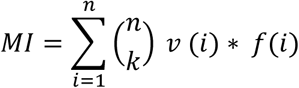

with *v(i)* being the c-p-value of a taxon i and *f(i)* being the frequency of that taxon in a sample.

Furthermore, nematode taxa were assigned to functional guilds according to Ferris et al. (2001)^50^, which then served as a basis to calculate additional indices. Functional guilds refer to the following trophic groups: bacterial feeders (Ba_X_), fungal feeders (Fu_x_), omnivores (Om_X_), and carnivores (Ca_X_). Associated numbers (i.e., the x of the respective trophic group) are again referring to the c-p values described above. The Enrichment Index (EI) indicates the responsiveness of the opportunistic bacterial (Ba_1_ and Ba_2_) and fungal feeders (Fu_2_) to food web enrichment^50^ and is calculated as follows:

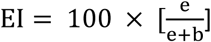

with *e* as weighted frequencies of Ba1 and Fu2 and *b* as weighted frequencies of Ba2 and Fu2 nematodes^50^. The Channel Index (CI) reflects the nature of decomposition channels through the soil food web. High values indicate a predominant decomposition pathway of organic matter dominated by fungal-feeding nematodes, whereas low values refer to bacterial-dominated decomposition pathways^50^.

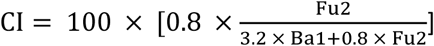

with 0.8 and 3.2 representing enrichment weightings for Fu_2_ and Ba_1_ nematodes^50^. The Structure Index (SI) provides information about the complexity of the soil food web. A highly structured food web with a high SI suggests ecosystem stability, while low values imply environmental disturbance^50^.

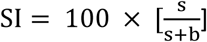

with *s* calculated as the weighted frequencies of Ba_3_-Ba_4_, Fu_3_-Fu_4_, Ca_3_-Ca_5_ and Om_3_-Om5 nematodes, and *b* representing the weighted frequencies of Ba_2_ and Fu_2_ nematodes^50^.

By plotting the Enrichment Index (EI) against the Structure Index (SI) we obtained a faunal profile that indicates, whether the nematode community can be described as basal and stressed or as structured, enriched and stable^50^.

## i. Statistical analyses

Linear mixed-effects models were used to analyse the effects of drought, NPK fertilization, season, and their interactions on invertebrate feeding activity, microbial activity, and microbial biomass using the R-package “*nlme*”^51^. The random intercept of the model was structured with plots nested within blocks, nested within year (year as a categorical factor). To account for repeated measurements within plots, we compared first-order autoregressive and compound symmetry covariance structures based on the Akaike information criterion (AIC). As differences between AIC values were lower than 2, the simplest covariance structure (i.e. compound symmetry) was used. Based on the importance of soil water content for microbial activity and biomass^52^, soil water content was added as an additional explanatory variable to the linear mixed-effects models (Tables S3-S4, Figs. S4-S5). As we were expecting a strong relation between aboveground plant biomass and microbial biomass^53^, additional linear mixed-effects models were used to test the influence of plant biomass on microbial biomass (Table S5, Fig. S6). Model assumptions were checked by visually inspecting residuals for homogeneity and Pearson residuals for normality. To meet the assumptions of the model, invertebrate feeding activity and microbial activity were log-transformed (log(x+1)). In addition, linear mixed-effects models were used to assess the effects of drought, NPK fertilization, season (spring and summer 2017), and their interactions on nematode indices, i.e. Enrichment Index, Structure Index, Channel Index, and Maturity Index. A random intercept with plots nested within block was included in the models. We accounted for repeated measurements within plots by using a compound symmetry covariance structure, which fitted the data better than a first-order autoregressive covariance structure based on the Akaike information criterion. To evaluate model variation explained by fixed and random effects, marginal and conditional R^2^ were calculated using the “*MuMIn*” package^54^; marginal R^2^ represents model variation explained by fixed effects in the final model and conditional R^2^ represents model variation explained by both random and fixed effects. Furthermore, generalized mixed-effects models (GLMM) were used to assess the effects of drought, NPK fertilization, season (spring and summer 2017), and their interactions on nematode richness, total density (i.e. total number of individuals in the nematodes community) and the abundance of each trophic group (i.e. percentage of individuals in each trophic group). Nematode richness and total density of nematodes were modelled with Poisson distribution, while the trophic groups were modelled with Binomial distribution. The random intercept of the model was structured with plots nested within blocks. To account for over-dispersion, an observation-level random effect was used in the model with omnivorous and predatory nematodes as a response variable. GLMM models were also used to assess the effects of drought, NPK fertilization, and their interactions on nematode functional guilds and cp-groups (Table S6) using Binomial distribution. The random intercept of the model was structured with plots nested within blocks, nested within sampling (sampling as a categorical factor). GLMM models were performed using the *“lme4”* package^55^. Figures are based on mixed-effects model fits extracted using the package *“ggeffects”*^*56*^. All statistical analyses were conducted using R version 3.4.2^57^.

## Results

### i. Soil microbial responses

Soil microbial activity ranged from 0.7 to 5.1 μl O_2_ h^−1^ g^−1^ dry weight soil with an average of 1.7 μl O_2_ h^−1^ per g dry weight soil across all measurements. We could not detect a significant effect of the drought or the fertilization treatment on soil microbial activity (Fig. 1a). However, microbial respiration was significantly affected by season, with lowest activity in summer and highest activity in winter (Table 1; Fig. 1b). In addition, we found a positive relation between microbial activity and soil water content (F_1,111_= 170.83, p < 0.001; Table S3) that was independent of the drought and fertilization treatment (Fig. S4).

**Table 1.** Results of linear mixed-effects models for the effects of drought, NPK fertilization, season, and their interactions on soil invertebrate feeding activity, soil microbial activity, and soil microbial biomass. A random intercept with plots nested within blocks, which were nested within year was added to the model. A compound symmetry covariance structure was used to account for repeated measurements within plots. Marginal R^2^: model variation explained by fixed effects; conditional R^2^: model variation explained by both fixed and random effects. Logarithmic transformations were used for soil invertebrate feeding activity and soil microbial activity. NPK = NPK fertilization. * p < 0.05; ** p < 0.01; *** p < 0.001.

| Response Variables | Drought | NPK | Season | Drought<br>x NPK | Drought x<br>Season | NPK x<br>Season | Drought x<br>NPK x<br>Season | $R^2$ % | |
| --- | --- | --- | --- | --- | --- | --- | --- | --- | --- |
|  | *F-value*<br>(1,27) | *F-value*<br>(1,27) | *F-value*<br>(3,108) | *F-value*<br>(1,27) | *F-value*<br>(3,108) | *F-value*<br>(3,108) | *F-value*<br>(3,108) | *marginal* | *conditional* |
| **Microbial activity** | 0.78 | 2.71 | 31.02*** | 0.02 | 1.07 | 0.30 | 0.28 | 37 | 49 |
| **Microbial biomass** | 2.25 | 35.48*** | 3.95* | 0.22 | 1.55 | 0.92 | 0.36 | 8.4 | 80 |
| **Invertebrate**<br>**feeding activity** | 22.65*** | 22.60*** | 22.78*** | 9.17** | 0.19 | 0.36 | 0.38 | 44 | 49 |

**Figure 1.**
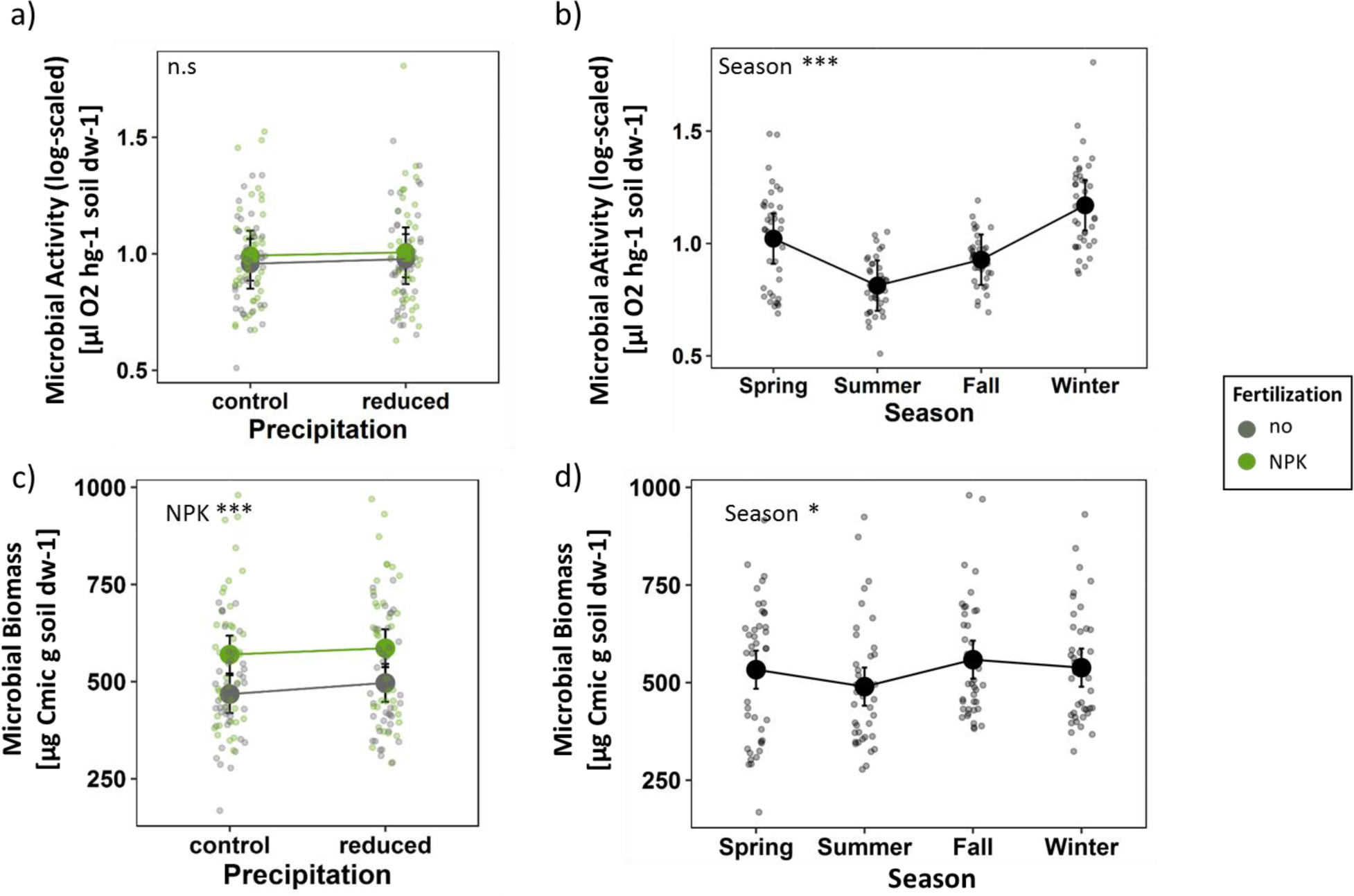
The effects of drought, fertilization, and season on soil microbial variables based on mixed effects model fits for each treatment. (a) Combined treatment effects across all seasons and (b) seasonal effects on soil microbial activity (log-scaled). (c) Combined treatment effects across all seasons and (d) seasonal effects on soil microbial biomass. Error bars indicate 95% confidence intervals. Grey = no NPK fertilization; green = NPK fertilization. n.s. = not significant; * p < 0.05; *** p < 0.001

Soil microbial biomass ranged from 168.0 to 979.8 μg Cmic g^−1^dry weight soil with an average of 530.6 μg Cmic g^−1^dry weight soil across all measurements. Overall, soil microbial biomass increased significantly with NPK fertilization (Fig. 1c). Microbial biomass was also significantly affected by season with lowest biomass in summer and highest biomass in fall (Fig. 1d, Table 1). Furthermore, fertilization and soil water content interactively affected microbial biomass (F_1,111_= 10.60, p = 0.002; Table S4); while microbial biomass increased with higher soil water content under ambient conditions, it slightly decreased with higher soil water content on plots with NPK fertilization (Fig. S5). In addition, soil microbial biomass increased significantly with aboveground plant biomass (F_1,59_= 8.81, p = 0.004; Table S6; Fig. S5).

### ii. Nematode responses

Neither total nematode density nor richness were significantly affected by any of the experimental treatments (Fig. 2a-b; Table 2); we could only detect significant differences between spring and summer (Fig. S7a-b). Among the nematode trophic groups, only bacteria-feeders were significantly increased by the fertilization treatment (Fig. 2c), which was mainly due to a significant increase of the r-strategic Ba_1_-nematodes (χ^2^ = 4.57, p = 0.032; Fig. S8a). In addition, bacteria-feeders were highly abundant in summer (Fig. S7c). Plant-feeders were affected by the interactive effects of fertilization and season: while fertilization favoured plant-feeding nematodes in spring, it had a negative effect in summer (Fig. 2d and S7d). The combined group of omnivorous and predatory nematodes marginally significantly decreased under drought and NPK fertilization (Fig. 2e), with a stronger negative effect of fertilization in spring (Fig. S7e). Fungal-feeders were not significantly affected by any of the treatments (Fig. 2f). A closer look at changes in the nematode community composition revealed that cp2 plant-feeding nematodes increased with drought (χ^2^ = 6.65, p = 0.0099; Fig. S9c; Table S6), whereas nematodes with higher c-p values, in detail Fu3-nematodes (χ^2^ = 4.97, p = 0.026; Fig. S8e), Om4-nematodes (marginally; χ^2^ = 3.80, p = 0.051; Fig. S8g), cp3 (marginally; χ^2^ = 2.74, p = 0.098; Fig. S9d), and cp4-nematodes (χ^2^ = 7.83, p = 0.0051; Fig. S9f) decreased significantly in response to fertilization.

**Figure 2.**
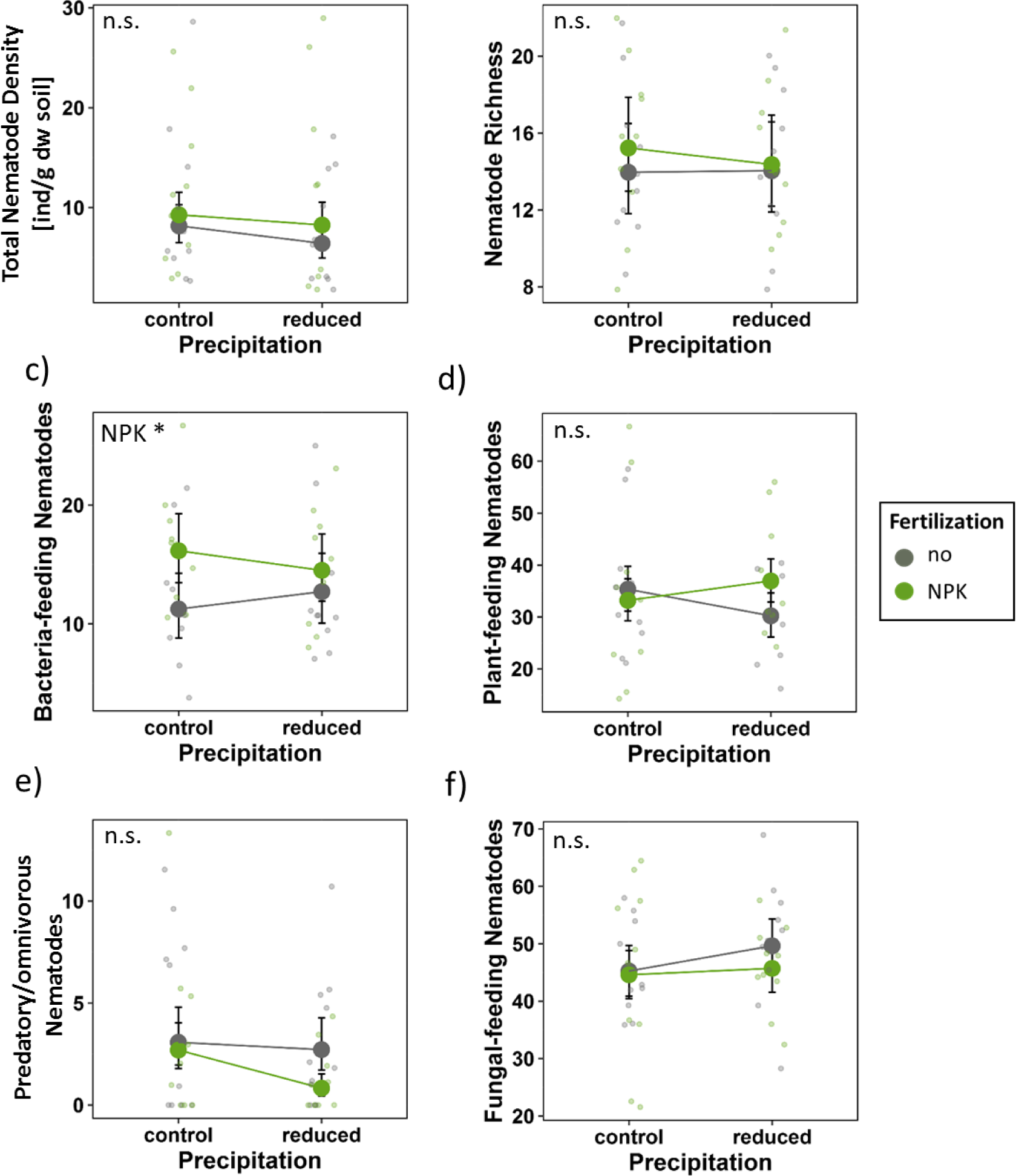
The effects of drought and fertilization on nematode response variables based on mixed effects model fits for each treatment. (a) Total nematode density per g dry weight soil; (b) nematode taxon richness; percentage of (c) bacteria-feeding nematodes; (d) plant-feeding nematodes; (e) fungal-feeding nematodes; and (f) predatory- and omnivorous nematodes. Error bars indicate 95% confidence intervals. Grey = no NPK fertilization; green = NPK fertilization. Both seasons (spring and summer 2017) are included (see Fig. S7 for seasonal effects). n.s. = not significant; * p < 0.05

**Table 2.** Chi-squared values (χ^2^) of the generalized mixed-effects models for the effects of drought, fertilization, season (spring and summer 2017), and their interaction on soil nematode density and richness using Poisson distribution and percentage of each nematode trophic group using Binomial distribution. Plots nested within blocks served as a random intercept in the model. NPK = annual NPK fertilization. (^*^) p < 0.1; * p < 0.05; *** p < 0.001

| Nematode response variable | Drought | NPK | Season | Drought x NPK | Drought x Season | NPK x Season | Drought x NPK x Season |
| --- | --- | --- | --- | --- | --- | --- | --- |
| | $\chi^2$ | $\chi^2$ | $\chi^2$ | $\chi^2$ | $\chi^2$ | $\chi^2$ | $\chi^2$ |
| Total density | 0.51 | 2.51 | 118.15*** | 0.66 | 0.80 | 0.47 | 0.79 |
| Richness | 0.14 | 0.38 | 21.47*** | 0.13 | 0.05 | 0.06 | 0.01 |
| Plant feeders | 0.24 | 1.22 | 12.53*** | 0.53 | 3.24<sup>(*)</sup> | 11.17*** | 1.10 |
| Fungal feeders | 1.15 | 0.74 | 0.57 | 0.06 | 3.24<sup>(*)</sup> | 1.87 | 0.61 |
| Bacteria feeders | 0.26 | 4.76* | 15.64*** | 0.71 | 2.05 | 0.25 | 0.95 |
| Predators/Omnivores | 3.53<sup>(*)</sup> | 3.50<sup>(*)</sup> | 0.80 | 2.16 | 1.53 | 5.63* | 0.07 |

**Table 3.** Results of linear mixed effects models for the effects of drought, fertilization, season (spring and summer 2017), and their interaction on soil nematode indices. Plots nested within blocks served as a random intercept in the model. A compound symmetry covariance structure was used to account for repeated measurements within plots. Marginal R^2^: model variation explained by fixed effects; conditional R^2^: model variation explained by both fixed and random effects. NPK = annual NPK fertilization. (^*^) p < 0.1; * p < 0.05; ** p < 0.01; *** p < 0.001

| Nematode response variable | Drought | NPK | Season | Drought x NPK | Drought x Season | NPK x Season | Drought x NPK x Season | R<sup>2</sup> % |  |
| --- | --- | --- | --- | --- | --- | --- | --- | --- | --- |
|  | *F-value*<br>(1,12) | *F-value*<br>(1,12) | *F-value*<br>(1,16) | *F-value*<br>(1,12) | *F-value*<br>(1,16) | *F-value*<br>(1,16) | *F-value*<br>(1,16) | *marginal* | *conditional* |
| Enrichment Index | 3.30 <sup>(*)</sup> | 0.84 | 0.07 | 0.00 | 0.03 | 0.05 | 0.17 | 9.4 | 18 |
| Structure Index | 4.95 <sup>(*)</sup> | 6.65* | 0.02 | 0.32 | 1.70 | 0.51 | 0.53 | 23 | 29 |
| Channel Index | 1.60 | 3.98 <sup>(*)</sup> | 1.42 | 0.04 | 1.62 | 0.00 | 0.11 | 19 | 50 |
| Maturity Index | 5.07 <sup>(*)</sup> | 10.18** | 0.00 | 0.52 | 1.99 | 0.21 | 0.72 | 28 | 34 |

While the Enrichment Index increased marginally significantly under drought conditions (Table 2; Fig. 3a), the Structure Index and the Maturity Index were marginally significantly decreased by drought (Fig. 3b-c). Fertilization increased the importance of the bacterial decomposition channel as indicated by a marginally significant decrease of the Channel Index (Fig. 3d). In addition, fertilization decreased nematode complexity as indicated by a significant decrease of the Structure Index and Maturity Index. (Fig. 3b-c). Furthermore, plotting the EI-SI profile of the nematode community grouped by the different treatment combinations depicted that while some of the control and drought plots could be found in quadrant C (undisturbed, moderate enrichment, structured food web), nearly all NPK and NPK x drought plots were located in quadrant D (stressed, depleted, degraded food web) or in quadrant A (high disturbed, enriched, disturbed food web) (Fig. 3e).

**Figure 3.**
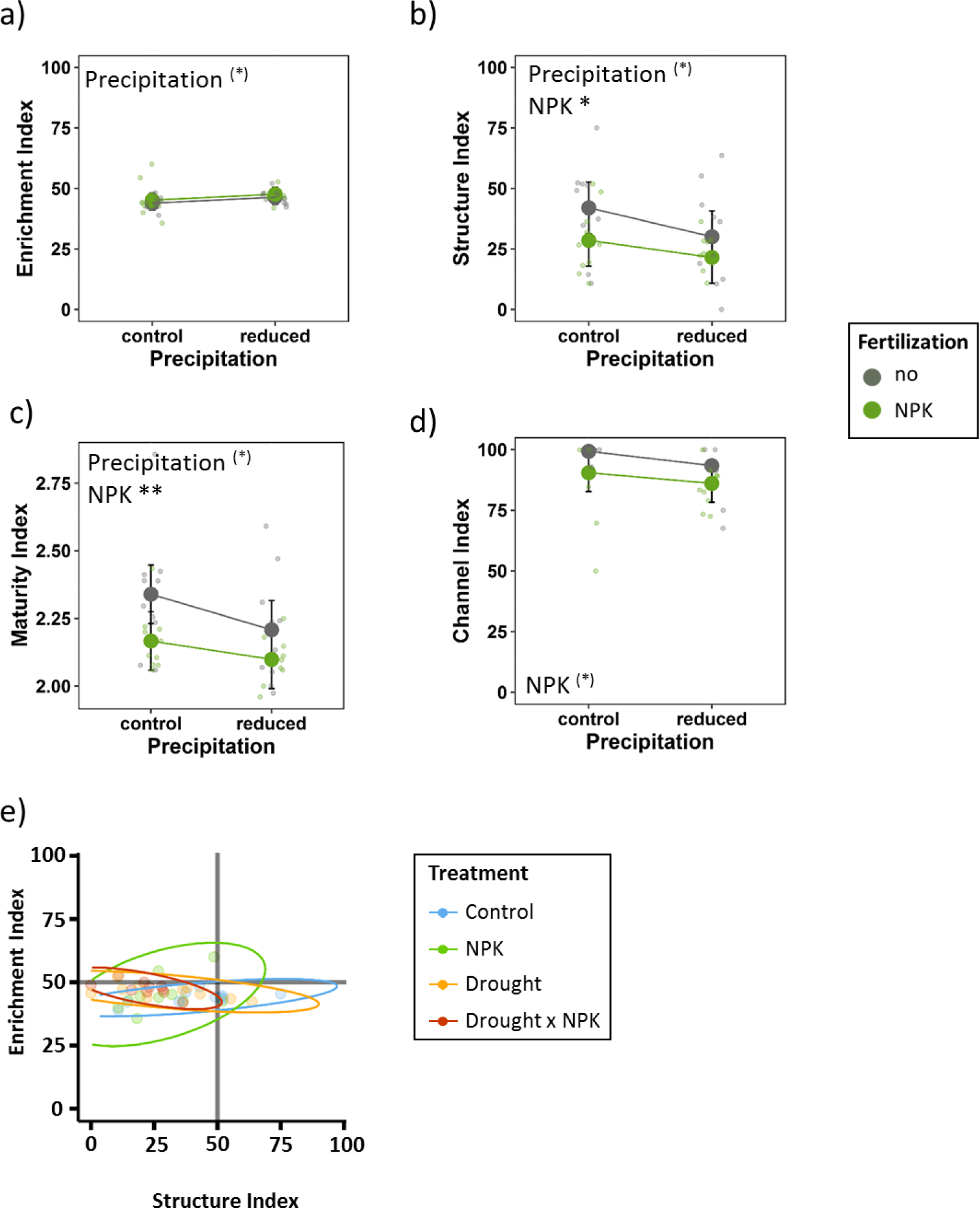
The effects of drought and fertilization on soil nematode indices based on mixed effects model fits for each treatment. (a) Enrichment Index; (b) Structure Index; (c) Maturity Index, and (d) Channel Index. Error bars indicate 95% confidence intervals. Grey = no NPK fertilization; green = NPK fertilization. (e) Enrichment Index (EI) and Structure Index (SI) trajectories for all plots according to the treatment combinations. Blue = control; green = NPK fertilization; yellow = drought; red = drought x NPK. Ellipses group treatments. Two seasons (spring and summer 2017) were included. ^(^*^)^ p < 0.1; * p < 0.05

### iii. Soil invertebrate feeding responses

Mean soil invertebrate feeding activity per plot ranged from 0 to 60% of consumed bait substrate with an average of 11% bait consumption. Feeding activity was significantly affected by an interactive effect of drought x fertilization; overall, fertilization decreased invertebrate feeding activity at ambient precipitation, but had no significant effect under drought conditions (Fig. 4a). These treatment effects were consistent across seasons (no treatment x season interaction), however, the level of soil invertebrate feeding activity varied strongly between seasons and tended to be highest in summer and strongly decreased in winter (Table 1; Fig. 4b).

**Figure 4.**
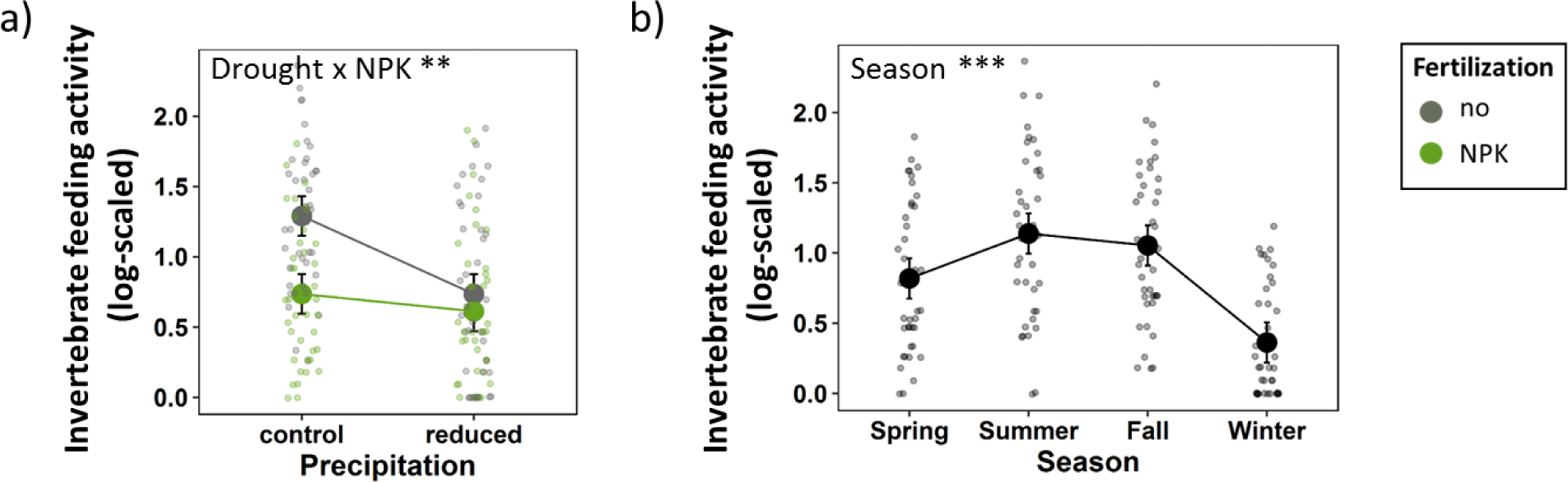
The effects of drought, NPK fertilization, and season on soil invertebrate feeding activity (log-scaled) based on mixed-effects model fits for each treatment. (a) Combined treatment effects across all seasons and (b) seasonal effects. Error bars indicate 95% confidence intervals. Grey = no NPK fertilization; green = NPK fertilization. ** p < 0.01; *** p < 0.001

## Discussion

We studied the impacts of two major global change drivers, namely drought and fertilization, and their interactive effects on soil communities and functions. By investigating the responses of a wide range of soil organisms across all seasons within a two-year timeframe, we gained a comprehensive picture of how soil ecosystem functions may be altered in a changing world. Intriguingly, we saw vastly different responses among trophic levels that were constant across seasons.

Our first hypothesis, predicting detrimental effects of drought on soil organisms, was confirmed to some extent: drought reduced soil invertebrate feeding activity and led to a more disturbed soil nematode community structure, whereas soil microbial activity and biomass were not significantly affected. The dependency of soil invertebrates on soil moisture content is well documented^58^ as is the fact that soil microfauna is more prone to water stress than bacteria and fungi^59^. The detrimental drought effects on faunal activity are thus in line with previous studies, which claim that abiotic conditions shape the performance of the soil faunal community^60^. Microarthropods (mites and collembolans), enchytraeids, and earthworms are some of the most relevant groups found in the upper soil layer in temperate regions^37^ and are likely to account for most of the feeding activity leading to bait perforation^61,62^. Extreme drought not only forces them to migrate to deeper soil layers, but also interferes with their reproduction and development success, which is possibly the reason why they are highly susceptible to drought^63-66^. Furthermore, drought conditions entail drier food sources for detritivores, which are more difficult to digest^41^.

In line with our expectations, drought was also responsible for (moderate) shifts in nematode indices. The environmental conditions were changed towards an enriched, more disturbed, less structured system, with a higher proportion of opportunistic nematodes. Already at ambient precipitation levels, the guilds of the opportunistic Fu_2_- and Ba_2_-nematodes accounted for the highest shares at our experimental site, indicating a basal food web that is capable to cover a wide ecological amplitude and is already adapted to some environmental stress^50^. The positive response of the Enrichment Index to drought suggests mortality at higher trophic levels, which subsequently promoted nutrient enrichment and gave further rise to opportunistic nematodes^50^. Thus, overall, drought led to simplified trophic structures of the nematode community.

In contrast to our initial hypothesis, soil microbial activity and biomass were not affected by the drought treatment. This was unexpected given the intensity of the drought treatment applied in the experiment (precipitation was reduced by 55% during the entire study period) and the fact that most soil microbes are strongly dependent on high soil moisture levels^52,67^ However, our results indicate that drought may not be a strong determinant of soil microbial activity and biomass at the study site. This is in line with Pailler et al. (2014)^68^, who found microbial functional responses to be robust against drought. In spite of the negligible responses to the drought treatment, we could reveal that soil water content explained a significant proportion of the variability in microbial activity, yet irrespective of the treatments. This suggests that microorganisms residing in the upper soil layer are highly depending on soil moisture levels and must therefore be able to sustain drought periods, for instance, through physiological modifications. Such adaptations may comprise an adjustment of internal water potential, sporulation, or production of exopolysaccharides that provide protection against exsiccation^69,70^ Apart from the resilience of the microbial community against experimental drought, we observed distinct variation of microbial activity and biomass across seasons. This provides evidence that temporal environmental variability is a strong predictor of species activity and emphasizes the dependency of soil organisms on seasonal patterns ^71-73^ We therefore infer that seasonal fluctuations in natural precipitation may have led to acclimatization of the microbial communities to drought periods as they are part of their climatic history ^74,75^ which may explain the weak effects of the experimentally induced drought.

The responses of soil biota were again ambiguous with regard to our second hypothesis, in which we expected fertilization to promote the activity of soil biota. Consistent with our hypothesis, soil microbial biomass increased under fertilization. Soil invertebrate feeding activity, however, substantially declined under elevated nutrient supply, questioning the universal validity of our initial hypothesis. The pronounced responsiveness of soil microbial biomass to NPK fertilization is in line with similar studies reporting positive effects of nitrogen fertilization on microbial biomass and changes in microbial community structure and function ^76,77^ Fertilization certainly enhanced nutrient availability, resulting in higher yields of aboveground plant biomass (Berger et al., in prep.), which is often accompanied by an expanded root system^28^. Subsequently, this increases the release of organic compounds into the soil^78^, providing substrates that support the growth of microbial communities towards higher population densities^53^.

Also nematode indices were highly responsive to NPK fertilization, resulting in a less structured and more disturbed system with an enhanced bacteria-driven decomposition pathway. The latter was additionally supported by a strong increase in bacteria-feeding nematodes (especially Ba1) and an equally strong decrease in fungal-feeding nematodes (Fu3) under fertilization, indicating that fertilization had detrimental effects on soil fungi. This is in line with the general notion that fertilization favours bacteria-dominated decomposition^18,27,50,79^. The simplification of the nutrient-enriched soil food web is also reflected by declines of the omnivorous nematodes (Om4) and the cp3 and cp4 nematodes, which consist of long-living, pollutant sensitive, rather immobile organisms that are prone to environmental stress^49^. When linking these changes reported for the nematode indices to the responses of the microbial community, our results suggest that fertilization promoted the growth of bacteria (simultaneously repressing fungi), which then accounted for a strong increase in microbial biomass. As a result, this restructuring of the microbial community may thus have provided the basis for the observed increase in bacteria-feeding nematodes and some of the shifts in nematode indices.

Current evidence for soil invertebrate responses to fertilization is equivocal. Several studies reported an enhancing effect of fertilization on soil invertebrate activity and diversity^80,81^, whereas others revealed no such effects^16,82^, or recorded declines in soil fauna abundances and diversity after fertilizer application^37^. Most likely, the diminished feeding activity of soil invertebrates can be explained by alterations of soil physicochemical properties. Fertilization often results in reduced soil pH, which is negatively correlated with the abundance of most soil invertebrates^83,84^. Especially earthworms, which represent a significant part of soil invertebrate biomass^85^, have been reported to decline noticeably in numbers under reduced soil pH^86^. In addition to this direct effect, the observed shifts in nematode indices could provide evidence for possible indirect effects on invertebrate feeding activity: as fungi seemed to be substantially reduced by fertilization, microarthropods like Collembola and Oribatid mites, which are strongly depending on fungi, may thereby be deprived of their main food source^82,87^. As a consequence, soil microarthropods may have declined in their abundances, thus limiting their possible contribution to bait perforation. Overall, environmental constraints, such as reduced soil pH in combination with altered energy channels and simplified food web structures, outweighed potential positive bottom-up effects through enhanced plant growth, which we expected to find.

Prior studies emphasized the importance of interactive effects of global change drivers as they will profoundly alter ecosystem functions and services^7,88^. Accordingly, our third hypothesis predicted fertilization to reinforce drought effects, resulting in reduced soil water content and thus aggravated living conditions for soil organisms. Indeed, drought and fertilization interacted significantly in restraining soil invertebrate feeding activity, partially supporting our hypothesis. However, as discussed above, fertilization obviously created an unfavourable environment for most soil invertebrates at our site. Although both global change drivers individually decreased soil invertebrate feeding activity, the interaction, however, did not result in further declines. We therefore conclude that the combined effects may have led to a distinct restructuring of the soil faunal community by promoting species better adapted to adverse conditions, and thus revealing no measurable change in net effects. Since the combined global change drivers are likely to modify aboveground plant communities^89-91^, alterations in the quality of aboveground litter inputs can be expected. Leaf litter quality, in turn, affects soil fauna and might therefore be responsible for reshaping the faunal community^92,93^. With the methods applied in our study, however, we can only speculate about potential changes in the soil faunal community composition. This highlights the need for future research to detect which specific groups are responsible for bait perforation. This could be done, for instance, by exposing bait lamina strips with a labelled substrate under controlled laboratory conditions^94,95^. Building on that, the abundances of the most important groups of soil organisms could be monitored in the field, while being exposed to different global change drivers.

In contrast to the invertebrate feeding activity, microbial activity was not significantly affected by the interaction of the two global change drivers. Moreover, we could not detect any interactive effects on nematode indices or nematode groups. This illustrates the robustness of a large portion of the soil community to interactive global change effects, which might therefore be able to buffer prospective global change effects to a certain extent.

In conclusion, the main groups of soil organisms investigated in the present study responded differently to the main and interacting effects of global change drivers. Soil invertebrate activity was strongly impaired by both global change drivers and their interaction, while microbial biomass benefited from enhanced nutrient availability, and microbial activity was surprisingly unaffected by all treatments. Despite the strong seasonal dynamics of temperate regions, these treatment effects remained constant across all seasons within two years. Notably, nematode indices pointed to changes in the state of the ecosystem, shifting towards simplified and more disturbed systems under drought and especially under fertilization that mostly facilitated opportunistic species. We could show that soil biota differ considerably in their sensitivity to global change drivers and in their seasonal dynamics - also highlighting the importance of integrating seasonal effects into experimental frameworks. This may lead to far-reaching alterations of crucial ecosystem processes, since decomposition and nutrient cycling are driven by the interdependent concurrence of soil microbial and faunal activities^40^. By covering a range of different taxonomic and trophic levels of soil organisms, we could therefore show that single as well interacting global change drivers induce complex changes in soil food webs and functions.

## Acknowledgements

We thank the staff of the Bad Lauchstädt Experimental Research Station for their help in maintaining the experimental site, and Alla Kavtea, Tom Künne, and Ulrich Pruschitzki for their support with lab and field work. Furthermore, we thank the coordination of the International Drought-Net Experiment for providing protocols and support. Financial support came from the German Centre for Integrative Biodiversity Research Halle-Jena-Leipzig, funded by the German Research Foundation (FZT 118).

